# Predictors of genomic diversity within North American squamates

**DOI:** 10.1101/2022.07.20.500845

**Authors:** Ivy E. Larkin, Edward A. Myers, Bryan C. Carstens, Lisa N. Barrow

**Author notes:** Current Address: Museum of Southwestern Biology and Department of Biology, University of New Mexico, Albuquerque NM 87131.

## Abstract

Comparisons of intraspecific genetic diversity across species can reveal the roles of geography, ecology, and life history in shaping biodiversity. The wide availability of mitochondrial DNA (mtDNA) sequences in open-access databases makes this marker practical for conducting analyses across several species in a common framework, but patterns may not be representative of overall species diversity. Here, we gather new and existing mtDNA sequences and genome-wide nuclear data (genotyping-by-sequencing; GBS) for 30 North American squamate species sampled in the Southeastern and Southwestern United States. We estimated mtDNA nucleotide diversity for two mtDNA genes, COI (22 species alignments; average 16 sequences) and cytb (22 species; average 58 sequences), as well as nuclear heterozygosity and nucleotide diversity from GBS data for 118 individuals (30 species; four individuals and 6,820–44,309 loci per species). We showed that nuclear genomic diversity estimates were highly consistent across individuals for some species, while other species showed large differences depending on the locality sampled. Range size was positively correlated with both cytb diversity (Phylogenetically Independent Contrasts: R^2^ = 0.31, p = 0.007) and GBS diversity (R^2^ = 0.21; p = 0.006), while other predictors differed across the top models for each dataset. Mitochondrial and nuclear diversity estimates were not correlated within species, although sampling differences in the data available made these datasets difficult to compare. Further study of mtDNA and nuclear diversity sampled across species’ ranges is needed to evaluate the roles of geography and life history in structuring diversity across a variety of taxonomic groups.

## Introduction

Genetic information is key for investigating the evolutionary histories and demographic trajectories of populations and species. Since the advent of DNA sequencing in the late 1970s (Sanger et al. 1977), analysis of sequence variation has enabled biologists to discover species, determine relationships among lineages, reveal demographic histories, and provide assessments of population structure and diversity (Avise 2000; Hebert et al. 2003; Brito and Edwards 2009). Genetic diversity within species is especially critical for persistence in changing environments and is clearly of interest for global conservation (Laikre 2010; Laikre et al. 2020), but several challenges remain for gathering and comparing genetic information from large numbers of species. The extent to which genetic diversity is predictable given species characteristics such as range size or life history traits is an area of active interest (e.g., Leffler et al. 2012; Romiguier et al. 2014; Singhal et al. 2017; Pelletier and Carstens 2018; Barrow et al. 2021). These studies demonstrate that predictors often differ across different taxonomic groups and scales, prompting the need for additional investigations into understudied clades.

The type of genetic data chosen for comparative studies is important to consider. Most work to date has used widely available markers from organellar genomes, but the extent to which these data are representative of broader genomic patterns remains unclear. Mitochondrial DNA (mtDNA) is easily sequenced and has been used for decades, making it readily accessible from public databases for comparisons across large numbers of species (Zink and Barrowclough 2008; Miraldo et al. 2016). While certain characteristics of mtDNA, such as a high mutation rate and lack of recombination, make it desirable for population-level studies and the inference of phylogenetic relationships at shallow scales (Avise et al. 1987), mtDNA may be poorly suited as a representation of genomic diversity in a given species (Galtier et al. 2009). As a nonrecombinant unit that is matrilineally inherited, mtDNA represents a single realization of many possible coalescent histories of sampled taxa, and it is well appreciated that any single genealogy may not accurately reflect population or species history (Hudson and Turelli 2003) particularly in cases where sex-biased dispersal or multiple divergences in a short timespan have occurred (Morin et al. 2004; Galtier et al. 2009). Notably, several investigations comparing genetic diversity from mitochondrial and nuclear genomes within species have found conflicting results across different taxonomic groups (Bazin 2006; Mulligan et al. 2006; Singhal et al. 2017b). Bazin et al. (2006) showed that mtDNA did not correspond with expectations of population abundance when comparing across broad animal groups, e.g., invertebrate groups did not have higher mtDNA diversity than vertebrate groups, while nuclear sequence and allozyme datasets did meet this expectation. Comparisons within vertebrate groups such as mammals (Mulligan et al. 2006) and lizards (Singhal et al. 2017) have demonstrated a positive correlation between estimates of mtDNA and nuclear diversity, however, suggesting that mtDNA may be a useful marker for understanding patterns of genetic diversity in animals with smaller population sizes. Taken together, the contrasting results of these studies also illustrate how the taxonomic scale of a given study can lead to different findings.

Sampling many nuclear loci enables more robust estimates of species relationships, population history, diversity estimates, and demographic parameters of interest (Edwards and Beerli 2000; Carling and Brumfield 2007). The growing availability of genome-scale nuclear datasets is promising for comparative studies (Garrick et al. 2015), but sampling many individuals and populations for multiple species in a single study remains cost prohibitive. One solution implemented in previous comparative studies is to obtain genome-wide estimates of diversity from a single or few individuals as a representative of each species (e.g., Romiguier et al. 2014; Singhal et al. 2017; Grundler et al. 2019). It is not yet clear how consistent genomic diversity estimates are across populations of wide-ranging species that have experienced different histories across their ranges (but see Nazareno et al. (2017) for an example in plant populations separated by 20 km). We begin to address this question in squamates by evaluating genomic diversity from multiple individuals and localities within species in the present study.

In addition to considering appropriate sampling strategies, identifying potential predictors of genomic diversity within species is challenging because many species-level characteristics may correspond with diversity within populations, diversity between populations (i.e., genetic structure), or both. Species with larger census population sizes are expected to have larger effective population sizes and therefore higher levels of neutral genetic diversity within populations (Kimura 1979). This assumed relationship may also extend to total range size, which has been considered as a proxy for census population size (e.g., Leffler et al. 2012; Singhal et al. 2017) because species that occupy large areas are presumed to be more locally abundant. The abundance-range size relationship may be explained by a variety of mechanisms (Gaston et al. 1997). In analyses of some taxonomic groups, population density appears to be unrelated to range size (Novosolov et al. 2017), while empirical examples of island and mainland bird species have demonstrated the expected positive relationship between genomic diversity and range size (Brüniche-Olsen et al. 2019; Leroy et al. 2021). Furthermore, the size, shape, and characteristics of a species’ range can impact how individuals move across the landscape, thus influencing rates of dispersal, gene flow, and the maintenance of genetic structure and overall diversity within species (Wright 1943; Sexton et al. 2014; Myers et al. 2019).

Life history and ecological traits also correspond with genetic diversity across broad taxonomic groups (Romiguier et al. 2014; Chen et al. 2017). Body size is negatively correlated with genetic diversity within and between populations for some groups (e.g., mammals: Brüniche-Olsen et al. 2018; bees: López-Uribe et al. 2019; butterflies: Mackintosh et al. 2019), which could relate to limits on population size (leading to lower within-population diversity) or to higher dispersal (leading to lower genetic structure) in larger-bodied species (White et al. 2007; Paz et al. 2015). Fecundity, or clutch size, is also expected to relate to abundance, where species that have more offspring will have larger and more stable population sizes through time and therefore higher neutral diversity (Kimura 1979). Indeed, clutch size is positively correlated with genetic diversity across animals (Romiguier et al. 2014), where species that have many, small offspring have higher within-population genetic diversity than long-lived species with few, large offspring. Aspects of mating system or reproductive mode have also been associated with genetic variation in different groups. In plants, within-population genomic diversity is higher in outcrossing species compared to selfing species (Chen et al. 2017), which is expected because inbreeding reduces effective population size.

Different reproductive strategies may also lead to differences in dispersal distance and resulting genetic structure. For example, in Panamanian frogs, species with direct development exhibit greater genetic structure compared to larval developing species (Paz et al. 2015); and in marine invertebrates, benthic direct-developing species have higher genetic structure than those with a pelagic larval stage (Collin 2001; Lee and Boulding 2009). Another aspect of reproduction that has not been thoroughly addressed is parity mode, whether species lay eggs (oviparous) or have live young (viviparous). Oviparity could be associated with higher dispersal, and less genetic structure, if females must travel long distances to locate suitable nest sites, while viviparous species have reduced movements because of increased energetic costs (Shine 2015). In the common lizard (*Zootoca vivipara*), greater genetic structure was found in an oviparous lineage compared to a viviparous lineage, but in this case, the pattern could be explained by differences in demographic history (Cornetti et al. 2015). Comparisons of species-wide genetic diversity across multiple oviparous and viviparous species are lacking thus far.

### North American Squamates in Regions of Contrasting Topographic Complexity

This study focuses on the genetic diversity of squamates from North America in two regions that were climatically suitable during the Last Glacial Maximum (LGM; Waltari et al. 2007), but differ in their topographic complexity (Badgley et al. 2017). We investigated species from the North American Coastal Plain and the Desert Southwest (Fig. 1), both of which harbor high reptile diversity (Jenkins et al. 2015) and are currently under threat from habitat fragmentation, climate change, and change in wildfire regimes (Archer and Predick 2008; Noss et al. 2015; Briggs et al. 2020). The North American Coastal Plain includes the Gulf and Atlantic coasts in the southeastern United States and has been described as a biodiversity hotspot (Noss et al. 2015). Although the region was never glaciated, when climatic and habitat shifts during Pleistocene glacial cycles occurred (Williams et al. 2004), the region likely provided stable refugial areas for many of the species in this study (Soltis et al. 2006; Weinell and Austin 2017). Within the Desert Southwest, located in the southwestern United States and north-central Mexico, the Chihuahuan Desert is the most biologically diverse desert and is considered among the 200 most biologically valuable ecoregions globally (Olson and Dinerstein 1998; Briggs et al. 2020). This region is also predicted to have provided refugial areas during the LGM (Waltari et al. 2007), but is more topographically complex with high environmental heterogeneity that plays an important role in population genetic structure and lineage divergence (Badgley et al. 2017; Myers et al. 2019). Both regions should retain much of the ancestral genetic diversity of the sampled species since each is suspected to have harbored refugial populations during the Pleistocene. Furthermore, the high squamate diversity in these regions provides the opportunity to compare species with a variety of range sizes, body sizes, and life history traits.

**Figure 1.**
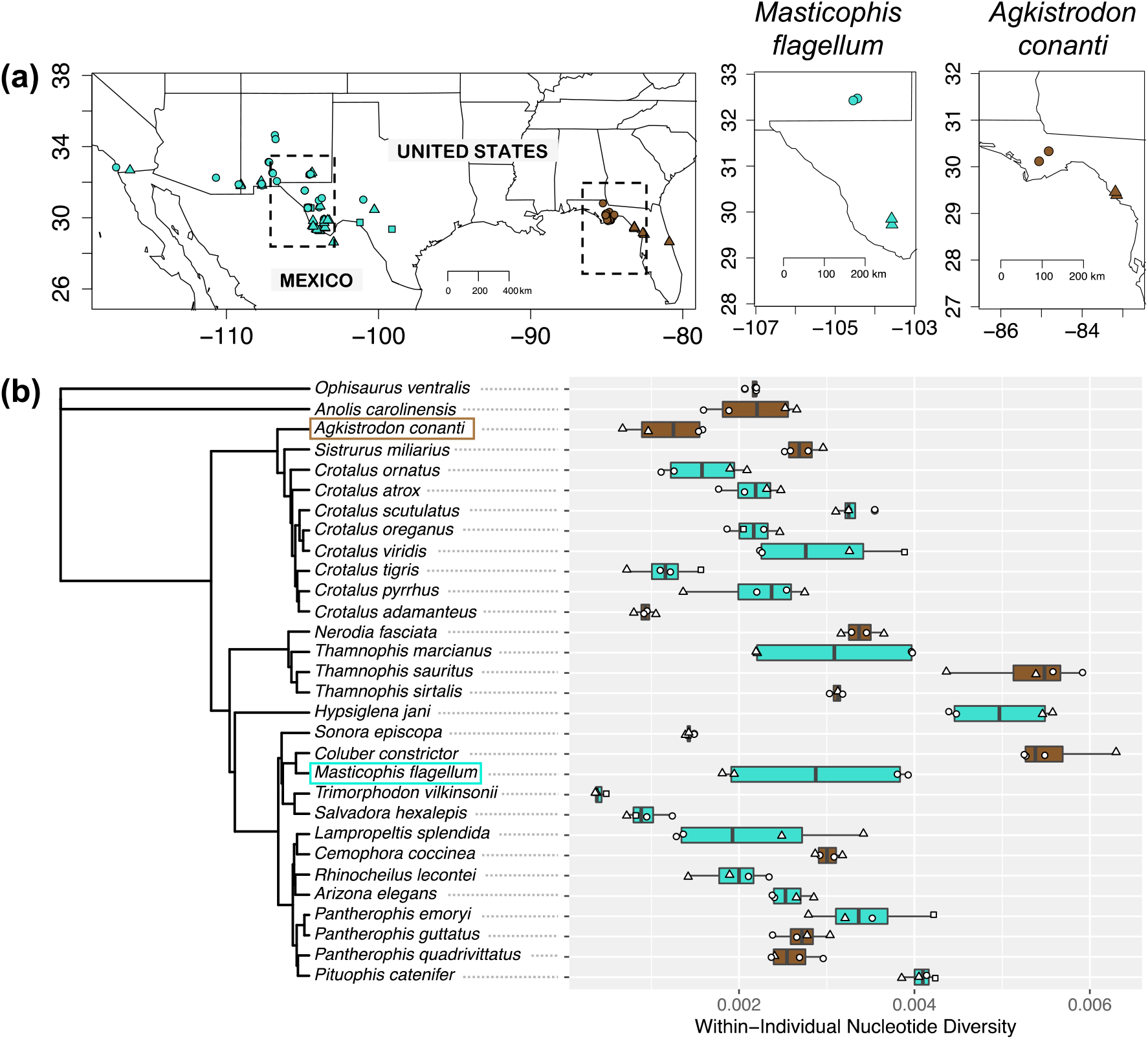
Individual sample localities and nucleotide diversity estimates for species included in the study. (a) Map of all samples for which GBS data were analyzed. Desert Southwest samples are shown in turquoise and Coastal Plain samples in brown. Dotted black boxes indicate the inset maps to the right, which depict localities sampled within two species as examples. Shapes indicate individuals were sampled from the same locality within a species. (b) Phylogenetic relationships for the 30 species included in the study based on Tonini et al. (2016). Within-individual nucleotide estimates are shown to the right with colors and shapes corresponding to the maps in (a).

In this study, we combined new and existing mtDNA and genome-scale nuclear sequence data for 30 North American squamate species to address three main questions. First, how consistent are nuclear genomic estimates across individuals and localities within species? Second, are measures of mtDNA and nuclear diversity within species correlated, suggesting that mtDNA is a useful proxy for within-species diversity? Third, are species geographic range size, body size, or life history traits (number of offspring and reproductive mode) associated with either mtDNA or nuclear diversity within squamates?

## Methods

### Sample Collection

We gathered and analyzed sequence data from 30 species representing four snake families (Viperidae, Colubridae, Natricidae, Dipsadidae) and two lizard families (Anolidae and Anguidae) (Fig. 1). We followed the most recent taxonomic revisions in the literature, ensuring mtDNA sequences could be assigned to a single species based on their specific localities, and excluding sequences without locality information (Table S1). Tissue samples for 12 Coastal Plain species (4–7 individuals each) were collected in northern Florida from 2009 to 2018, preserved in either 95% ethanol or DMSO tissue buffer, frozen at or below −20 °C, and subsequently archived at the Museum of Southwestern Biology (Fig. 1; Table S2). We also included previously published data for 18 snake species distributed across the Desert Southwest (Myers et al. 2019, 2020; Myers 2021). For the nuclear dataset of these 18 species, we chose four individuals with a similar sampling strategy as the Coastal Plain species. Briefly, we chose two individuals each from two localities per species when possible, focusing primarily on the Chihuahuan Desert and a similar distance (∼200 km) between localities. This standardized distance enabled us to make comparisons among species without the potentially confounding effects of geographic distance on genetic structure.

### Range Maps and Trait Data

Species range maps were downloaded as shapefiles from the IUCN Red List of Threatened Species Version 6.1 (www.iucnredlist.org/). Maps were edited in QGIS 3.12 as needed to exclude non-native parts of the range where species have been introduced and to edit species ranges according to recent taxonomic changes (Table S1). The R packages ‘rgdal’ (Bivand et al. 2021) and ‘geosphere’ (Karney 2013; Hijmans 2021) were used to visualize ranges and calculate total range area in kilometers squared. Species trait information was obtained primarily from Burbrink and Myers (2015), including maximum body size (log-transformed), average clutch size (log-transformed), and parity (viviparous or oviparous). Trait data for the two lizard species were obtained from field guides and online natural history accounts (Powell et al. 2016; https://www.virginiaherpetologicalsociety.com; https://animaldiversity.org).

### Mitochondrial Sequences

For the 12 Coastal Plain species, genomic DNA was extracted from tail or liver tissue with the E.Z.N.A. DNA Tissue Kit (Omega Bio-Tek) according to the manufacturer’s instructions. We amplified a 658-base pair portion of the mtDNA cytochrome c oxidase 1 gene (COI) using primers ReptBCF_COI and ReptBCR_COI (Castañeda and de Queiroz 2011). The thermal profile included an initial denaturation step at 94° C for 3 minutes, 35 cycles of denaturing at 94° for 45 seconds, annealing at 53° for 30 seconds, elongating at 72° for 60 seconds, and a final extension step at 72° for 10 minutes. PCR products were visualized on agarose gels to ensure successful amplification, cleaned with EXOSAP-IT (Affymetrix Inc.), and sequenced at The Ohio State University (OSU) Comprehensive Cancer Center Genomics Shared Resource. Species-specific sequencing primers were designed when the original PCR primers did not perform adequately (Table S3). Additional mtDNA sequences for two genes, COI and cytochrome b (cytb), were obtained from NCBI GenBank for species with data available. Sequences for each species were aligned in Geneious 2020.1.1 (https://www.geneious.com) using MAFFT 7.450 (Katoh and Standley 2013).

### Nuclear Data Collection

Genotyping-by-sequencing (GBS) libraries were prepared following a modified version of the protocol described in Elshire et al. (2011). For four samples per Coastal Plain species (48 individuals), we digested 100 ng of input DNA with the enzyme Pst1, ligated 8 μL of adapter mix including unique barcodes, pooled libraries, and performed a bead cleanup with Sera-Mag Speedbeads (Rohland and Reich 2012). Final PCR amplification was conducted with 16 cycles in 8 replicate reactions, followed by pooling, bead cleanup, and quantification via a Qubit fluorometer (Life Technologies). Libraries were size selected to 200-500 bp using a Blue Pippen (Sage Science Inc.), quantified with a Bioanalyzer (Agilent Technologies), and sequenced at the OSU Comprehensive Cancer Center Genomics Shared Resource on an Illumina HiSeq 4000 with 150 bp paired-end sequencing. For four samples per Desert Southwest species (72 individuals), we downloaded GBS reads from the NCBI Sequence Read Archive (Table S2).

Sequenced GBS reads from all 30 species were assembled through the ipyrad 0.7.28 pipeline (Eaton and Overcast 2020) with the following settings. After demultiplexing (no mismatches allowed in barcodes), the R1 reads for the four individuals per species were assembled to generate within-species datasets. We trimmed reads for the Southeastern U.S. species to 125 bp prior to assembly using the built-in ipyrad option which uses the software tool ‘cutadapt’ (trim_reads = 0, −25). We used the de novo assembly method with a clustering similarity threshold of 85% (clust_threshold = 0.85), maximum clustering depth of 10000 reads (maxdepth = 10000), maximum of five low quality base calls per read (max_low_qual_bases = 5; with Q<20), minimum depth for base calling of six reads (mindepth_statistical = 6 and mindepth_majrule = 6, strict filtering for adapters (filter_adapters = 2), and we retained reads longer than 35 bp after trimming adapters (filter_min_trim_len = 35). We allowed a maximum of eight heterozygous sites in consensus sequences (max_Hs_consens = 5), a maximum of 20 SNPs per locus (max_SNPs_locus = 20) and required all four individuals (no missing data) to be included in a locus to retain that locus in output files (min_samples_locus = 4).

### Genetic Diversity Metrics

Measures of mtDNA diversity were calculated for each species using the R package ‘pegas’ (Paradis 2010). We calculated nucleotide diversity (pi or π, Nei 1987) for COI and cytb alignments of each species with the nuc.div function. Nuclear diversity within species was estimated from each GBS dataset using the R packages ‘adegenet’ (Jombart 2008; Jombart and Ahmed 2011) and ‘PopGenome’ (Pfeifer et al. 2014). Expected heterozygosity (Hs) was computed from the unique SNPs outfile from ipyrad for each species using ‘adegenet’. To calculate π, ‘PopGenome’ reads in a directory of FASTA files including an alignment for each locus; we used a custom Python script (available on Dryad) to generate this directory for each species using the alleles outfiles (full sequences) from ipyrad. We calculated π within each sample (two alleles per individual) and then determined the mean and standard deviation across individuals, hereafter referred to as “mean within-individual π”. For comparison, we also calculated π within each species from the alignments including four individuals (eight alleles) per species, hereafter “within-species π”. Species-level and within-population nuclear diversity can be estimated from a single individual when sufficient loci are sampled (e.g., Chen et al. 2017; Grundler et al. 2019). Where possible, we included two individuals per species from each sampled locality to investigate how consistent genomic diversity estimates are across different individuals and localities within species.

### Statistical Analyses

We compared mtDNA (COI π, cytb π) and nuclear (mean within-individual π, within-species π) diversity estimates within species using pairwise correlations in R. We then constructed phylogenetic generalized least square (PGLS) models to assess the importance of different predictors on genetic diversity, while accounting for the possibility that closely related species may share similar genomic diversity and species traits. We downloaded 100 phylogeny subsets for the 30 species in our study from VertLife.org, which includes the squamate relationships from Tonini et al. (2016), and generated a least-squares consensus tree using phytools (Revell 2012) for our analyses. We constructed a model set for each of three responses: mtDNA COI π, mtDNA cytb π, and nuclear within-species π. For each response, predictors of interest included geographic range size, maximum body size, average clutch size, and parity. To assess effects of sample size on diversity metrics, we included the number of individuals sampled as a covariate in mtDNA models, and the number of GBS loci as a covariate for nuclear models. We included the region (Coastal Plain or Desert Southwest) in our initial PGLS models for nuclear within-species π and used Welch’s two-sample t-tests to check for differences between the GBS datasets generated in this study and those generated previously.

Analyses were conducted using the R packages ‘ape’ (Paradis and Schliep 2019), ‘geiger’ (Harmon et al. 2008), and ‘nlme’ (Pinheiro et al. 2022). We constructed PGLS models using two models of evolution, Brownian Motion (BM) and Ornstein-Uhlenbeck (OU). The BM model assumes traits evolve according to a random walk, while the OU model incorporates a trait optimum towards which traits are pulled (Felsenstein 1985; Martins and Hansen 1997). For each model set, all combinations of the predictors of interest were included and models were ranked based on the corrected Akaike Information Criterion (AICc). We ran model selection analyses using the R package ‘MuMIn’ (Bartón 2022) to determine which model of evolution had a better fit. We then constructed models incorporating all combinations of the predictors of interest and ran model selection analyses to assess which sets of predictors were included in the best-fit models for each diversity measure. We further assessed relationships between continuous variables using Spearman’s rank correlation tests, phylogenetically independent contrasts implemented in ‘ape’ (Felsenstein 1985), and linear models.

## Results

### Data Summary

We generated 62 new COI sequences (GenBank accession numbers ON911378– ON911439) and combined these with available sequences on GenBank. Of the 30 squamates in our study, we were able to analyze COI alignments for 22 species (4–69 sequences per alignment; average 16) and cytb alignments for 22 species (8–143 sequences; average 58). Three species (*Crotalus oreganus, Hypsiglena jani, Trimorphodon vilkinsonii*) did not have sufficient mtDNA data available for either COI or cytb and 17 species had alignments for both genes (Table 1). We sequenced new GBS data for 47 individuals from 12 species (NCBI SRA accession numbers SAMN31800422–SAMN31800468; PRJNA903383) and calculated genomic diversity metrics for 118 individuals from 30 species. One *Thamnophis sirtalis* sample had poor sequencing success and was removed from subsequent analyses. The assembled GBS datasets included 2,923–25,086 loci for the unique SNP datasets and 6,820–44,309 loci for the full sequence datasets depending on the species (Table 1).

**Table 1.** Sample sizes for mtDNA and GBS datasets for each species.

| Species | Family | # COI<br>seqs | # cyt b<br>seqs | # GBS<br>samples | # GBS loci<br>(unique<br>SNPs) | # GBS loci<br>(seqs) |
| --- | --- | --- | --- | --- | --- | --- |
| *Ophisaurus ventralis* | Anguidae | 6 | NA | 4 | 22951 | 42875 |
| *Anolis carolinensis* | Anolidae | 16 | NA | 4 | 15708 | 28909 |
| *Agkistrodon conanti* | Viperidae | 5 | 46 | 4 | 5961 | 15540 |
| *Sistrurus miliarius* | Viperidae | 10 | 31 | 4 | 14356 | 23527 |
| *Crotalus ornatus* | Viperidae | NA | 27 | 4 | 9047 | 29251 |
| *Crotalus atrox* | Viperidae | 5 | 66 | 4 | 6926 | 19307 |
| *Crotalus scutulatus* | Viperidae | 4 | 74 | 4 | 14696 | 29502 |
| *Crotalus oreganus* | Viperidae | NA | NA | 4 | 11032 | 27615 |
| *Crotalus viridis* | Viperidae | NA | 8 | 4 | 15565 | 29627 |
| *Crotalus tigris* | Viperidae | 5 | NA | 4 | 9003 | 37903 |
| *Crotalus pyrrhus* | Viperidae | 6 | NA | 4 | 3274 | 6820 |
| *Crotalus adamanteus* | Viperidae | 10 | 126 | 4 | 8424 | 27387 |
| *Nerodia fasciata* | Natricidae | 30 | 143 | 4 | 15130 | 25148 |
| *Thamnophis marcianus* | Natricidae | NA | 22 | 4 | 7467 | 17648 |
| *Thamnophis sauritus* | Natricidae | 7 | NA | 4 | 14540 | 17945 |
| *Thamnophis sirtalis* | Natricidae | 7 | 37 | 3 | 15664 | 25409 |
| *Hypsiglena jani* | Dipsadidae | NA | NA | 4 | 19304 | 29950 |
| *Sonora episcopa* | Colubridae | 7 | 20 | 4 | 2923 | 12181 |
| *Trimorphodon wilkinsonii* | Colubridae | NA | NA | 3 | 5566 | 26644 |
| *Salvadora hexalepis* | Colubridae | 8 | 31 | 4 | 3513 | 20475 |
| *Masticophis flagellum* | Colubridae | 55 | 136 | 4 | 7512 | 18350 |
| *Coluber constrictor* | Colubridae | 51 | 22 | 4 | 18737 | 22267 |
| *Lampropeltis splendida* | Colubridae | NA | 25 | 4 | 6485 | 19095 |
| *Cemophora coccinea* | Colubridae | 69 | 9 | 4 | 15437 | 25907 |
| *Rhinocheilus lecontei* | Colubridae | 9 | 105 | 4 | 4713 | 15022 |
| *Arizona elegans* | Colubridae | 8 | 92 | 4 | 16879 | 37596 |
| *Pantherophis quadrivittatus* | Colubridae | 12 | 20 | 4 | 14352 | 26136 |
| *Pituophis catenifer* | Colubridae | 9 | 98 | 4 | 18205 | 32161 |
| *Pantherophis emoryi* | Colubridae | NA | 129 | 4 | 25086 | 44309 |
| *Pantherophis guttatus* | Colubridae | 11 | 11 | 4 | 14377 | 25517 |

### Mitochondrial and Nuclear Diversity

Mitochondrial diversity within species ranged from 0.0015 to 0.0812 for COI π and from 0.0005 to 0.0717 for cytb π. Diversity estimates for the two mtDNA genes were correlated within species (Fig. S1; Spearman’s rho (ρ) = 0.897, p < 2.2e-16). For the nuclear GBS data, within-species heterozygosity ranged from 0.3 to 0.346 and within-species π ranged from 0.0005 to 0.0083. Mean within-individual π ranged from 0.0004 to 0.0056 and there was substantial variation among individuals and localities for some species (Fig. 1). For species such as *Thamnophis marcianus* and *Masticophis flagellum*, the within-individual π estimates were consistent between the two individuals within a locality, but values for one locality were nearly double the values for the second locality. For other species such as *Crotalus adamanteus* and *Sonora episcopa*, the within-individual π estimates were highly consistent for all four individuals. For still other species such as *Thamnophis sauritus* and *Lampropeltis splendida*, there was large variation in π estimates even for individuals collected from the same locality.

The within-species π estimates were strongly correlated with the mean within-individual π estimates (Fig. S1; Spearman’s ρ = 0.986, p < 2.2e-16), and we used the within-species estimate to represent overall species nuclear diversity for subsequent analyses. Metrics of diversity for mtDNA and nuclear GBS data were not correlated for the species represented in both datasets (Fig. 2). Nuclear within-species π was not correlated with mtDNA COI π (Spearman’s ρ = 0.16, p = 0.476) or mtDNA cytb π (Spearman’s ρ = 0.293, p = 0.185). We did not detect any clear differences in nuclear diversity between datasets sampled from the Desert Southwest compared to the Coastal Plain (Fig. S2).

**Figure 2.**
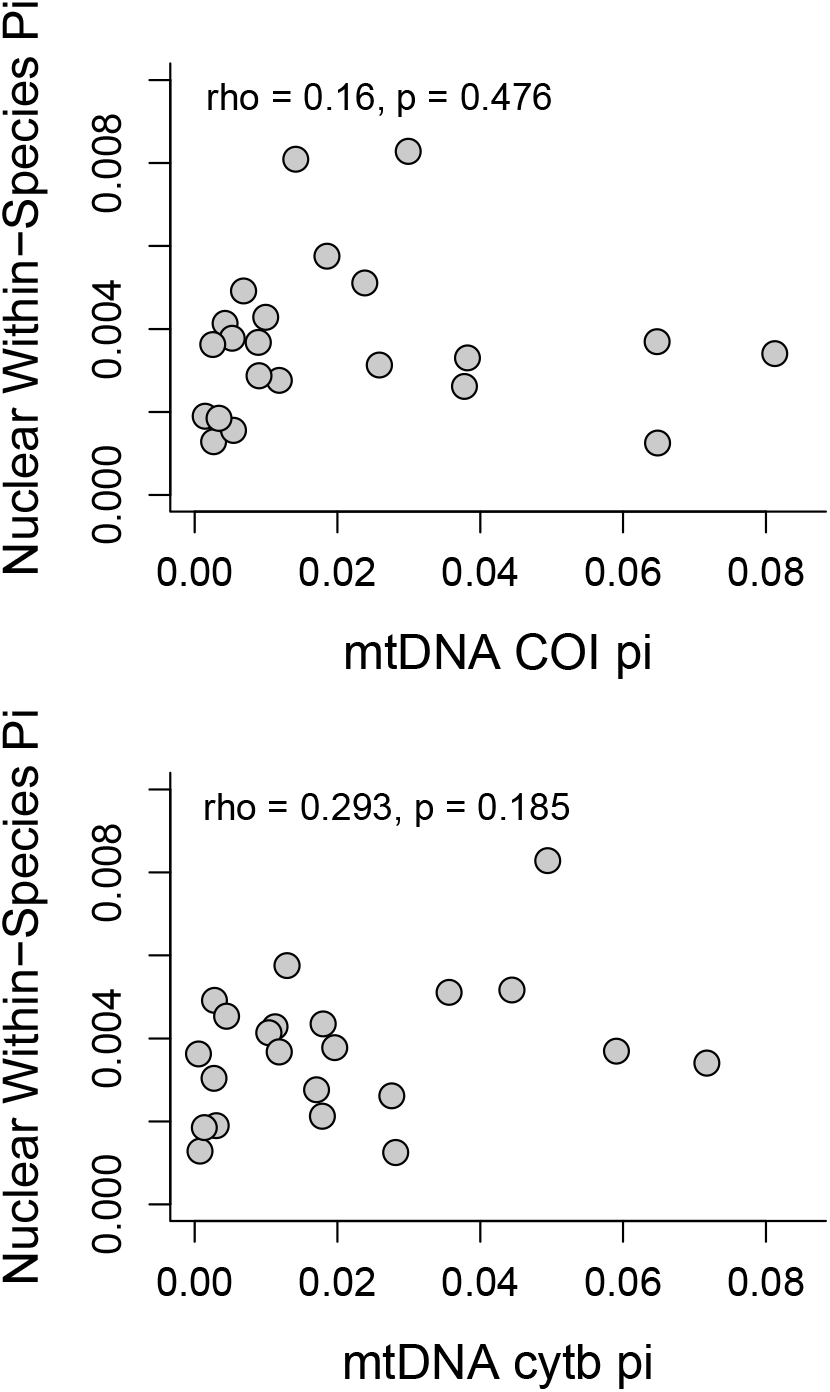
Lack of correlation between mtDNA and nuclear diversity estimates within species. Spearman’s rank correlation sample estimates and p-values are shown.

### Predictors of Squamate Genetic Diversity

The OU model of evolution was the better fit for all three response variables (model weights > 0.986; Table 2). Different sets of predictors were included in the top models for the three response variables (Table 3). For mtDNA COI, parity was the only variable consistently included in the top models. Diversity between viviparous and oviparous species was not significantly different according to the top PGLS model (Table 4), although oviparous species did have higher COI diversity than viviparous species without phylogenetic correction (Fig. 3a; t-test: t = 2.34, p = 0.03). For mtDNA cytb, range size and parity were always included in the top models. Within-species cytb diversity had a positive relationship with range size (Fig. 3), suggesting that species with larger ranges tend to have higher mtDNA diversity. This relationship was consistent for both uncorrected (Spearman’s ρ = 0.64; p = 0.002; Fig. 3c) and phylogenetically corrected datasets (phylogenetically independent contrasts: R^2 = 0.31; p = 0.007; Fig. 3d). Oviparous species had higher cytb diversity compared to viviparous species (Fig. 3b; t-test: t = 3.11, p = 0.008).

**Table 2.** Model comparison of PGLS models under the Brownian Motion (BM) or Ornstein-Uhlenbeck (OU) models of evolution.

| Response | Model | log Likelihood | AIC | ΔAIC | Weight |
| --- | --- | --- | --- | --- | --- |
| mtDNA COI $\pi$ | OU | 57.452 | -98.9 | 0.00 | 0.998 |
|  | BM | 50.320 | -86.6 | 12.26 | 0.002 |
| mtDNA cytb $\pi$ | OU | 66.965 | -117.9 | 0.00 | 0.997 |
|  | BM | 60.257 | -106.5 | 11.42 | 0.003 |
| Within-species $\pi$ | OU | 157.485 | -297.0 | 0.00 | 0.994 |
|  | BM | 151.435 | -286.9 | 10.1 | 0.006 |

**Table 3.**
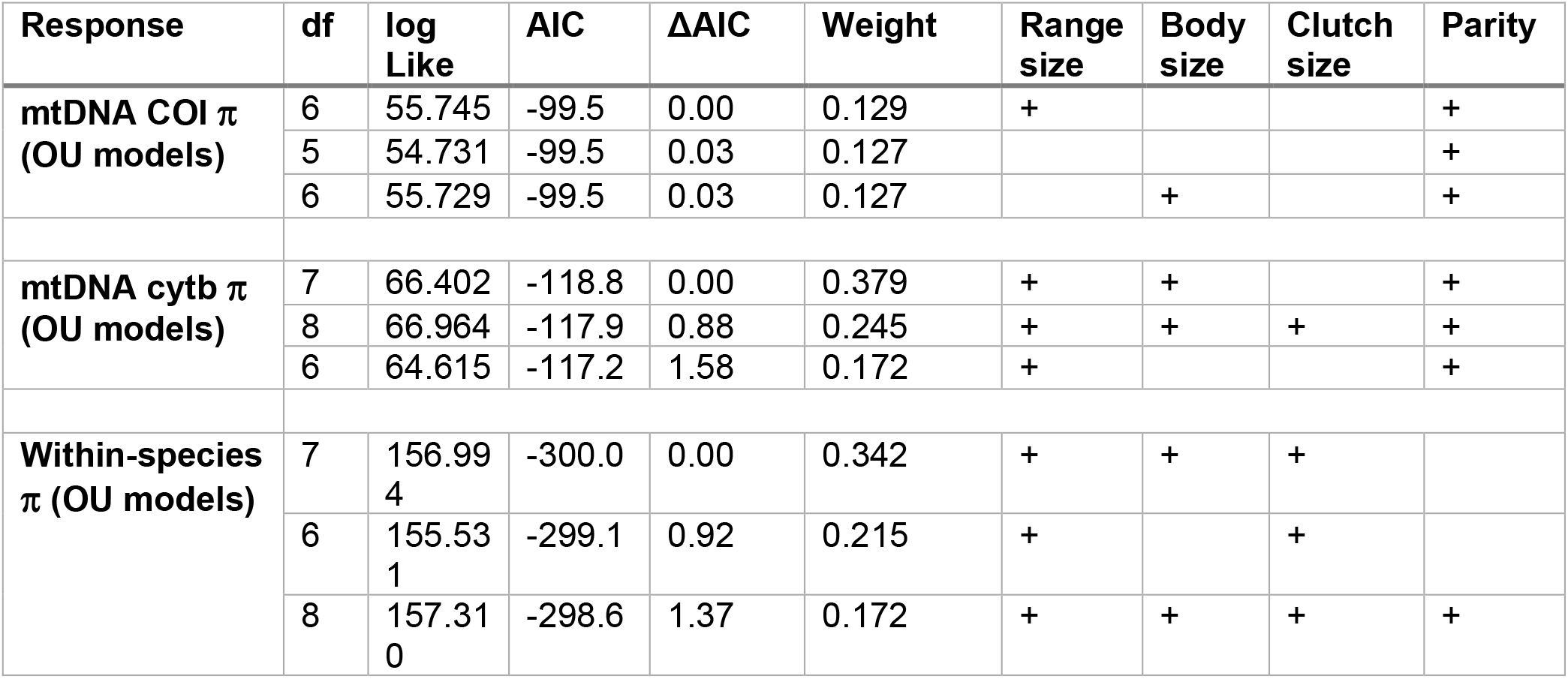
Model comparison of PGLS models with different sets of predictors. The top three models for each response are shown, which all had a delta AIC value of <= 2. Note that none of the models for mtDNA COI π were substantially improved over others (delta AIC all < 4.4). The plus (“+”) sign in the predictor columns indicates that predictor was included in that model.

**Table 4.** Model coefficients, standard errors (SE), t, and p-values for the top model listed in Table 3 for each response.

| Response | Coefficient | Value | SE | t | p-value |
| --- | --- | --- | --- | --- | --- |
| **mtDNA COI $\pi$**<br>**(OU model)** | Intercept | -0.0565 | 0.0662 | -0.8540 | 0.4043 |
|  | log(Range size) | 0.0062 | 0.0047 | 1.3184 | 0.2039 |
|  | Parity_vivip | -0.0165 | 0.0100 | -1.6530 | 0.1157 |
|  | N sequences | -0.0001 | 0.0003 | -0.2777 | 0.7844 |
| **mtDNA cytb $\pi$**<br>**(OU model)** | Intercept | -0.1522 | 0.0539 | -2.8252 | 0.0117 |
|  | log(Range size) | 0.0078 | 0.0033 | 2.3328 | 0.0322* |
|  | log(Body size) | 0.0315 | 0.0182 | 1.7320 | 0.1014 |
|  | Parity_vivip | -0.0179 | 0.0061 | -2.9503 | 0.0090* |
|  | N sequences | 0.0001 | 0.0001 | 0.8222 | 0.4223 |
| **Within-species $\pi$**<br>**(OU model)** | Intercept | -0.0099 | 0.0041 | -2.4112 | 0.0236 |
|  | log(Range size) | 0.0010 | 0.0003 | 3.8158 | 0.0008* |
|  | log(Body size) | -0.0022 | 0.0014 | -1.6001 | 0.1221 |
|  | log(Clutch size) | 0.0043 | 0.0017 | 2.5383 | 0.0177* |
|  | N loci | 0.0000 | 0.0000 | 0.0665 | 0.9475 |

**Figure 3.**
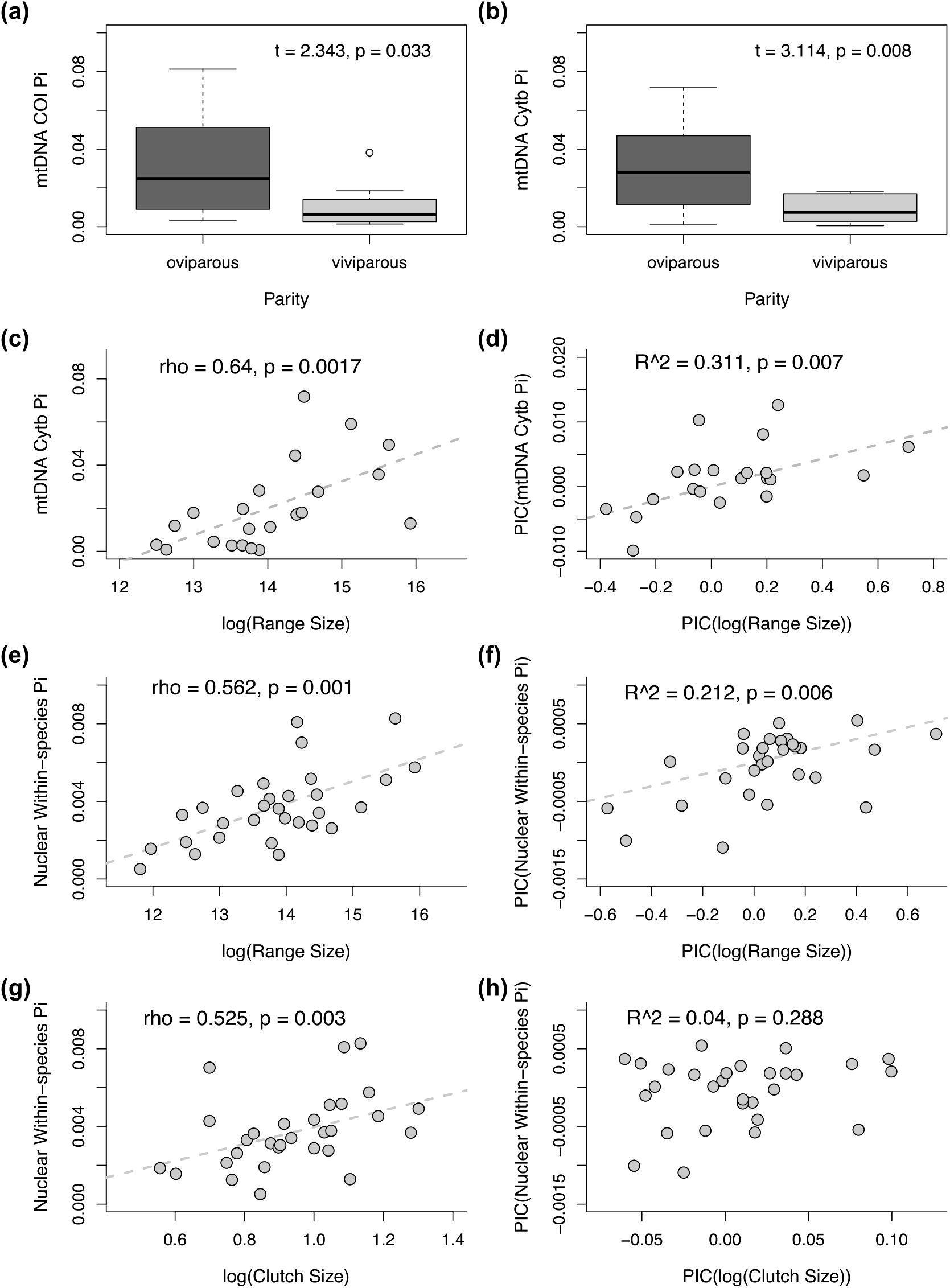
Predictors of mtDNA (a-d) and nuclear diversity (e-h) within species. a, b) Welch’s two-sample t-test statistics and p-values are shown. c, e, g) Scatter plots and dotted lines depict simple linear relationships without phylogenetic correction. Spearman’s rank correlation sample estimates and p-values are shown. d, f, h) Phylogenetically-corrected R^2^ values and p-values are shown.

Range size and clutch size were consistently included in the top models for nuclear within-species π (Table 2). Nuclear within-species π had a positive relationship with both range size and average clutch size (Table 4; Fig. 3). The relationship between nuclear π and range size was consistent for both uncorrected (Spearman’s ρ = 0.56; p = 0.001; Fig. 3e) and phylogenetically corrected datasets (PIC: R^2 = 0.21; p = 0.006; Fig. 3f). The relationship between nuclear π and average clutch size was significant for the uncorrected data (Spearman’s ρ = 0.53; p = 0.003; Fig. 3g), but not based on phylogenetically independent contrasts (R^2 = 0.04; p = 0.288; Fig. 3h).

## Discussion

We compared genetic diversity estimates from mitochondrial and nuclear genomes for 30 North American squamate species. Our analyses demonstrated that range size is positively correlated with mtDNA and nuclear diversity in squamates and that life history traits are inconsistent predictors of diversity for these datasets. We found that mtDNA and nuclear diversity estimates were not correlated, but we note that several aspects of sampling made these datasets difficult to compare. Here, we discuss sampling considerations for this and future studies, including the number of individuals, localities, species, and loci available or feasible to sample. Despite these challenges, we provide empirical evidence that geography is a key element for understanding patterns of genomic diversity within and across species.

### Sampling Considerations Across Geography, Taxonomy, and Genomes

Comparative population genomics is a developing field that aims to uncover the forces underlying genomic diversity by comparing across multiple species (Ellegren and Galtier 2016; Edwards et al. 2022). Studies typically focus on the roles of selection, neutrality, and recombination in structuring genetic variation in a primarily non-geographic context, thus sampling usually consists of a single or few individuals to represent each species (e.g. Romiguier et al. 2014; Chen et al. 2017). In contrast, comparative phylogeography takes a geography-centric view to understanding historical processes underlying current genetic variation within species and lineages. Ideally, species have been sampled across their entire range and as densely as possible, often revealing the presence of multiple lineages within nominal species (e.g., Schield et al. 2018). A third emerging field, macrogenetics, focuses on an even broader geographic and taxonomic scale, typically using repurposed data from relatively few genetic markers to investigate large-scale patterns and predictors of genetic variation within species (Leigh et al. 2021). Macrogenetic studies rely on sampling choices and data from previous studies, but until relatively recently, genome-wide datasets were not common in phylogeographic studies (Garrick et al. 2015). The gradual accumulation of these data over time and the introduction of new data aggregators (e.g. Pelletier et al. 2022) potentially increases the scale of comparative studies that use opportunistic sampling, but the novel data collected here offer a direct comparison between two ecoregions.

Thoroughly sampling many individuals from many species across geographic space remains costly, thus a pressing question relevant to the design of comparative studies is: How representative are genomic diversity estimates from a few individuals or from a narrow part of a species’ range? Our GBS data indicate that this answer depends on several factors – certain species (e.g., *Crotalus adamanteus*) had highly consistent genomic diversity estimates from localities sampled ∼200 km apart, while others (e.g., *Thamnophis marcianus*) had two-fold differences in genomic diversity between localities (Fig. 1). This result indicates that using a single individual or locality to represent diversity for an entire species does not always provide a complete picture of diversity within species. We expected to find similar levels of genetic diversity across individuals because we sampled them from historically stable regions. Thus, it may seem surprising that such differences are already emerging. We predict even greater differences when populations are sampled from topographically complex regions or in areas that have been influenced by historical climate change (e.g., Howes and Lougheed 2008; Nali et al. 2020), a consideration that should continue to be addressed in future studies.

Our study provides an initial comparison of squamate genomic diversity from five families sampled in two species-rich regions, the North American Desert Southwest and North American Coastal Plain. We found considerable variation in diversity among species within some families (e.g., Colubridae and Natricidae; Fig. 1), but further taxon sampling is needed for robust comparisons of genomic diversity across families. In general, the Desert Southwest has more heterogenous environments and landscapes compared to the Coastal Plain, and Florida in particular, where populations were sampled (Noss et al. 2015; Badgley et al. 2017). Thus far, we did not detect any clear differences in average genomic diversity levels between the two regions (Fig. S2), although several Desert Southwest species had high variation in genomic diversity among individuals and localities (Fig. 1). This result is expected given the heterogeneous nature of the region, where large differences in climate, elevation, and habitat on relatively small spatial scales can conceivably lead to differences in population sizes and standing genomic diversity among nearby localities (Myers et al. 2019).

We found that mtDNA and GBS diversity were not correlated, suggesting that mtDNA may not be a reliable proxy for within-species diversity in North American squamates. Several differences in our sampling for mtDNA and nuclear data made these datasets difficult to compare, however. The number of mtDNA sequences available for some species was limited, producing COI and cytb alignments for only 22 species each. These mtDNA sequences most likely represent a much broader geographic area within each species compared to our GBS data, though it is not possible to quantify the influence of geographic distance because sample localities were not associated with the majority of GenBank sequences (Marques et al. 2013). On the other hand, the GBS data consist of several thousand loci and provide a better representation of genome-wide diversity within species compared to a single mtDNA gene. These differences in power and geographic representation, along with relatively few data points for comparison (22 species), may explain the lack of correlation between mtDNA and nuclear diversity metrics in our study. Previous studies have found a relationship between mtDNA and nuclear diversity in some taxonomic groups including mammals (47 species; Mulligan et al. 2006) and Australian lizards (60 species; Singhal et al. 2017). Further study of this topic is important for comparative studies because mtDNA is readily available for many more species, but reduced representation nuclear genome datasets are increasing and have the potential to provide new insights on overall species diversity.

### Additional Challenges for Repurposed Data

One challenge for comparative studies such as ours is the difficulty of gathering different data types from species complexes that have undergone recent taxonomic revisions. When possible, we manually edited mtDNA alignments and IUCN range maps to reflect data from a single described lineage (Table S1). In some cases, however, recent taxonomic revisions and limited geographic information about existing sequences and traits made it impossible to assign data to newly-described species. For those taxa, we excluded mtDNA data and assumed trait information from former species designations would be representative for the whole complex. We provide the alignments and GenBank accession numbers for the data we analyzed (https://doi.org/10.5061/dryad.qfttdz0ks) such that they can be reassigned and reanalyzed in follow-up studies. Future efforts to link geographic coordinates with genetic sequences in open access databases will continue to improve the prospects of comparative studies (Pelletier et al. 2022). Frequent updates to geographic distribution and trait databases will also be needed to facilitate large-scale comparative studies.

The approaches used to obtain and analyze nuclear genome-wide datasets are important to consider for comparative studies of genetic diversity. One appealing characteristic of mtDNA data for data repurposing and comparing species sequenced across different studies is the ease of aligning known genes. Restriction site associated methods such as GBS provide many more loci for analysis, but without an annotated reference genome for most species of interest, these loci are anonymous. Furthermore, the locus assemblies and resulting parameters are highly dependent on the parameter settings of the pipeline used for analysis (Harvey et al. 2016; Shafer et al. 2017). Our study used existing squamate datasets from the Desert Southwest that we reassembled to ensure the same ipyrad version and parameters were used for comparison with new GBS data we collected for the Coastal Plain species. We found that heterozygosity values were highly sensitive to the parameter settings used. Prior to reassembly, heterozygosity estimates for the Desert Southwest species were an order of magnitude smaller than those of the Coastal Plain species (Table S4). These contrasting results demonstrate the additional challenges of comparing GBS and similar datasets collected from different studies. Targeted sequence capture approaches may be preferable to increase comparability across studies, though they remain more expensive and require more initial effort to design probe kits (Harvey et al. 2016; Singhal et al. 2017a; Hutter et al. 2022).

### Predictors of Squamate Genomic Diversity

Range size was positively correlated with nucleotide diversity within squamate species for both mtDNA (cytb) and nuclear data, despite differences in geographic and locus sampling. Previous studies have found mixed results in a variety of taxonomic groups and with different sampling strategies. For example, a global study including single-gene sequences from more than 8,000 eukaryotic species found that total range size was one of the most important predictors of genetic structure within species (Pelletier and Carstens 2018). In contrast, (Romiguier et al. 2014) analyzed transcriptome data from 2-10 individuals of 76 animal species and found no correlation between genomic diversity and range size. Detailed focus within taxonomic groups demonstrated correlations between range size and genetic diversity in *Drosophila* (Leffler et al. 2012), Australian lizards (Singhal et al. 2017b), and cetaceans (Vachon et al. 2018), but not in butterflies (Mackintosh et al. 2019). To our knowledge, we provide the first evidence that total range size may be a useful predictor of squamate genomic diversity. Future studies including additional species, biogeographic regions, and nuclear sampling across species’ ranges would provide important insights about the generality of these findings and the potential influence of fluctuations in population size on genomic variation.

The life history and ecological traits we included were less consistent predictors of mitochondrial and nuclear diversity in squamates. Body size has negative associations with genetic diversity in various taxonomic groups including mammals (Brüniche-Olsen et al. 2018), frogs (Paz et al. 2015), and insects (López-Uribe et al. 2019; Mackintosh et al. 2019). One possible explanation is that larger-bodied animals can disperse over longer distances, leading to less genetic structure and overall diversity across a species range. Our results do not provide compelling evidence for squamates, perhaps indicating that body size is not a good indicator of dispersal for this group. Our results for clutch size are consistent with the prediction that species with larger clutch sizes harbor higher genomic diversity (e.g., Romiguier et al. 2014), though this relationship did not remain after phylogenetic correction (Fig. 3h). These patterns will require further scrutiny with increased species sampling in future studies.

The potential association of parity with mitochondrial diversity, where oviparous (egg-laying) species had higher diversity on average than viviparous (live-bearing) species, is intriguing and should also be investigated further with more comprehensive sampling. Since the mitochondrial sequences were sampled across species ranges, our results may reflect greater genetic structure for oviparous species, in contrast with the prediction that oviparous species must disperse to find suitable nest sites (Shine 2015). The evolution of viviparity is associated with colder climates and faster development times in cool temperatures (Ma et al. 2018), but our study lacked sampling across broad latitudinal or climatic gradients. Furthermore, most of the oviparous species included in this study belong to the family Colubridae, but viviparity in squamates has evolved from oviparity at least 34 and perhaps more than 100 times (Pyron and Burbrink 2014; Blackburn 2015). Squamates therefore present an excellent opportunity to continue investigating whether life history traits such as clutch size or parity are associated with intraspecific genomic diversity while taking phylogeny into account.

Overall, we provide new insights into the predictors of diversity within squamates considering both mitochondrial and nuclear genome patterns. We showed that geographic range size corresponds with intraspecific genomic diversity even when a relatively small part of the range is sampled. We also demonstrate that genomic diversity can vary widely between populations on small spatial scales, suggesting that sampling a few individuals may not adequately represent diversity levels within species. As genome-wide datasets continue being generated, comparative studies of intraspecific diversity across species ranges will increase understanding of the roles of geography and life history in structuring diversity across a variety of taxonomic groups.

## Acknowledgements

We thank Moses Michelsohn for providing tissues from Florida and Megan Smith for laboratory and analytical support. Samples were obtained in accordance with collecting permits from the Florida Fish and Wildlife Conservation Commission and animal care protocols approved by the Florida State University Animal Care and Use Committee.

## Data Availability

We have deposited the primary data underlying these analyses as follows:

- Sampling locations, field numbers, museum catalogue numbers, R/Python scripts, and datasets used for analyses and visualization: Dryad link - https://doi.org/10.5061/dryad.qfttdz0ks
- DNA sequences: GenBank accession numbers ON911378–ON911439; NCBI SRA accession numbers: SAMN31800422–SAMN31800468 (PRJNA903383)

## Supplementary Materials

Table S1. Taxonomy used for this study and details about how different datasets were handled to reflect recent taxonomic changes. References that were used to manually edit mitochondrial alignments and range maps are provided.

*[provided as Excel sheet – tab 1]*

Table S2. List of all samples for which mtDNA COI was sequenced and GBS data were analyzed. Sample names, museum catalogue numbers, geographic coordinates, locality information, and total number of GBS reads sequenced are included.

*[provided as Excel sheet – tab 2]*

Table S3. Additional species-specific sequencing primers designed in this study to improve COI sequencing results.

*[provided as Excel sheet – tab 3]*

Table S4. Data for 30 squamate species used as input for analyses.

*[provided as Excel sheet – tab 4 and with PGLS R script in Dryad package]*

Table S5. Data for 118 squamate individuals used to make figures and summaries.

*[provided as Excel sheet – tab 5 and with R script in Dryad package]*

Additional files on Dryad (https://doi.org/10.5061/dryad.qfttdz0ks): Mitochondrial (COI and cytb) alignments including previous GenBank samples; GBS files; Range maps; Python/R scripts

**Figure S1.**
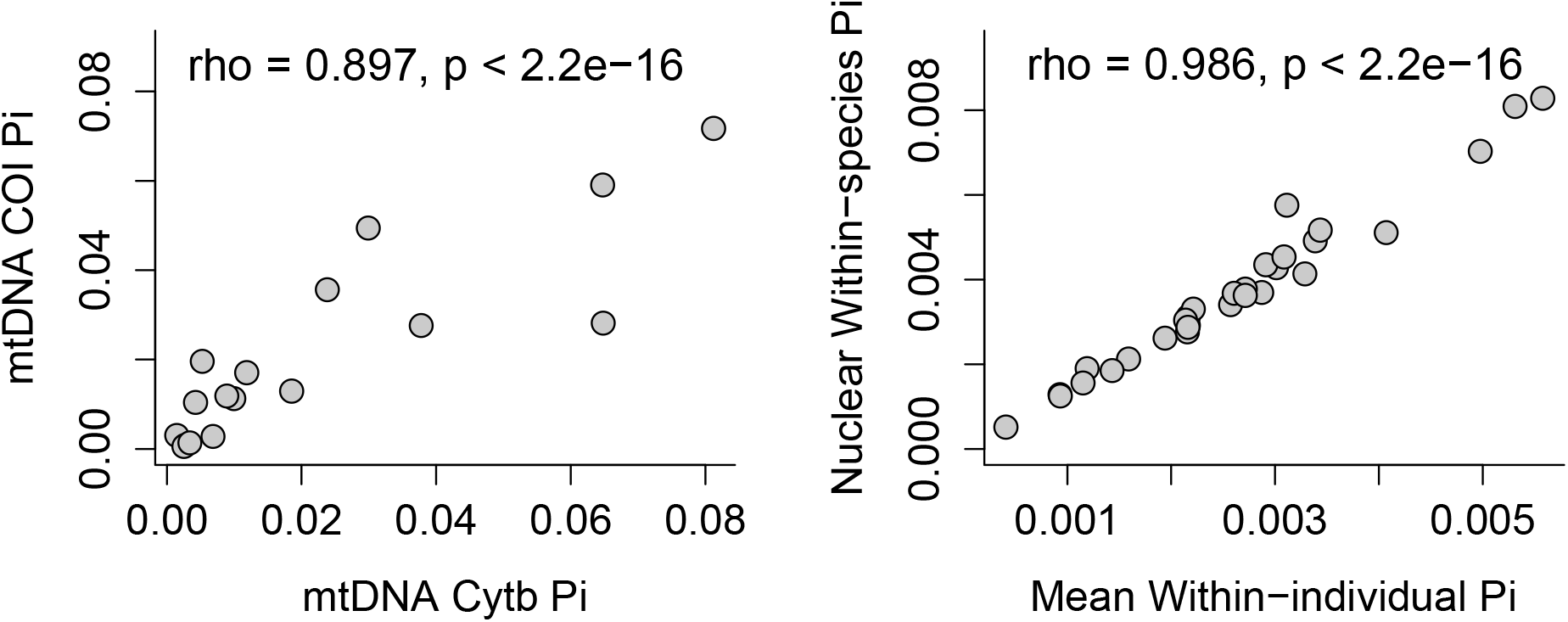
(a) Diversity estimates for mtDNA genes COI and cytb are correlated. (b) Mean within-individual pi estimates are strongly correlated with within-species pi estimates.

**Figure S2.**
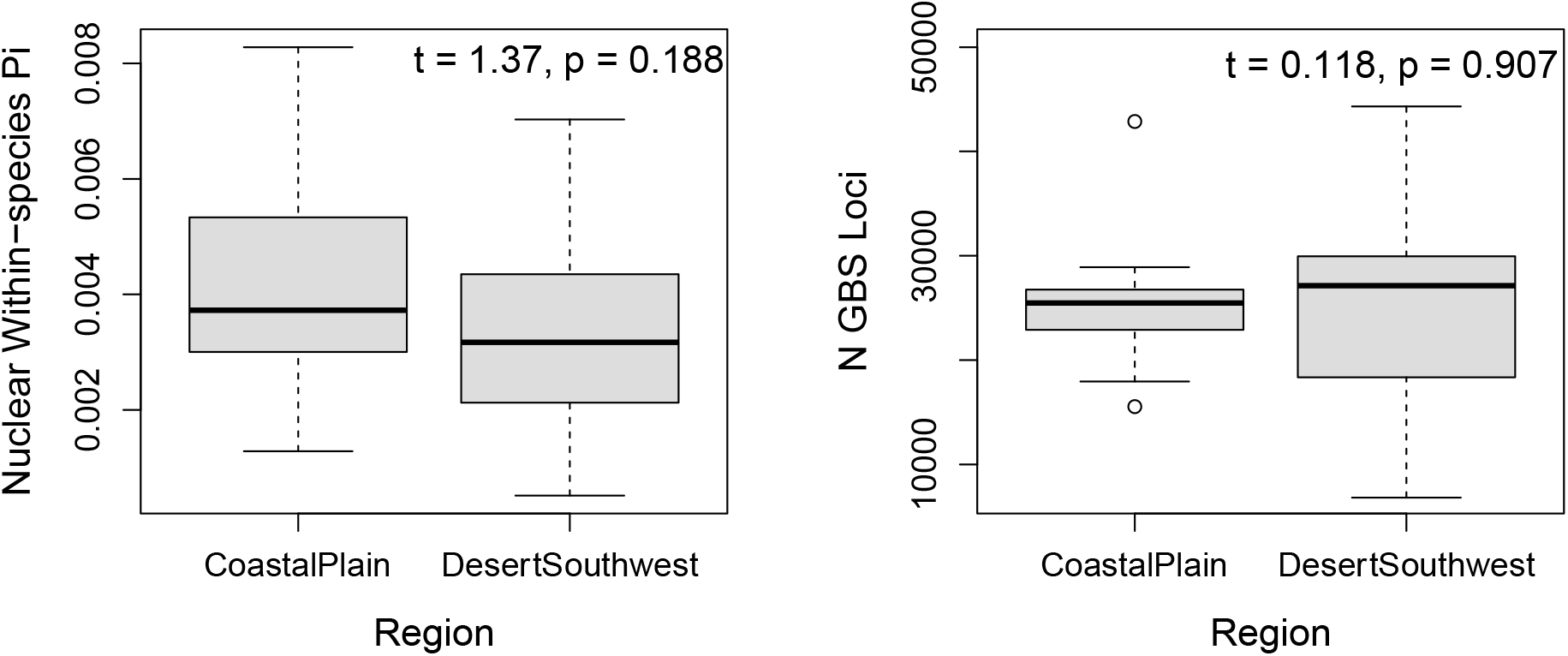
There are no clear differences in nuclear diversity estimates or the GBS datasets collected from the two different regions.

